# Antagonism of STAT3 signalling by Ebola virus

**DOI:** 10.1101/2020.08.10.245464

**Authors:** Angela R. Harrison, Megan Dearnley, Shawn Todd, Diane Green, Glenn A. Marsh, Gregory W. Moseley

## Abstract

Many viruses target signal transducers and activators of transcription (STAT) 1 and 2 to antagonise antiviral interferon (IFN) signalling, but targeting of signalling by other STATs/cytokines, including STAT3/interleukin (IL-) 6 that regulate processes important to Ebola virus (EBOV) haemorrhagic fever, is poorly defined. We report that EBOV potently inhibits STAT3 responses to IL-6 family cytokines, and that this is mediated by the IFN-antagonist VP24. Mechanistic analysis indicates that VP24 effects a unique strategy combining distinct karyopherin-dependent and karyopherin-independent mechanisms to antagonise STAT3-STAT1 heterodimers and STAT3 homodimers, respectively. This appears to reflect distinct mechanisms of nuclear trafficking of the STAT3 complexes, revealed for the first time by our analysis of VP24 function. These findings are consistent with major roles for global inhibition of STAT3 signalling in EBOV infection, and provide new insights into the molecular mechanisms of STAT3 nuclear trafficking, significant to pathogen-host interactions, cell physiology and pathologies such as cancer.

**Author summary:** Ebola virus (EBOV) continues to pose a significant risk to human health globally, causing ongoing disease outbreaks with case-fatality rates between 40 and 60%. Suppression of immune responses is a critical component of EBOV haemorrhagic fever, but understanding of EBOV impact on signalling by cytokines other than interferon is limited. We find that infectious EBOV inhibits interleukin-6 cytokine signalling *via* antagonism of STAT3. The antagonistic strategy uniquely combines two distinct mechanisms, which appear to reflect differing nuclear trafficking mechanisms of critical STAT3 complexes. This provides fundamental insights into the mechanisms of pathogenesis of a lethal virus, and biology of STAT3, a critical player in immunity, development, growth and cancer.

## Introduction

Outbreaks of Zaire ebolavirus (EBOV, family *Filoviridae*, order *Mononegavirales*) cause severe haemorrhagic fever with fatality rates between 40 and 60% [1–4]. The 2014-2016 West African outbreak (> 11,000 human deaths), and recent outbreak in the Democratic Republic of Congo (c. 2300 deaths in 2018-2020) highlight the ongoing danger to human health [3, 4].

The capacity of mammalian viruses to overcome the type-I IFN-mediated antiviral innate immune response is an important factor in virulence [5–7]. IFNs are induced in response to cellular detection of viral infection, and signal in autocrine and paracrine fashion to activate intracellular signalling, principally through STAT1 and STAT2. Following IFN-receptor binding, STAT1/2 are phosphorylated at conserved tyrosines, which results in the formation of phospho-(pY-)STAT1-STAT2 heterodimers and pY-STAT1 homodimers. Nuclear localisation signals (NLSs) formed within the dimers bind to nuclear import receptors of the NPI-1 karyopherin subfamily (which include karyopherin alpha-1 (Kα1)) at a ‘non-classical’ cargo-binding site, distinct from sites bound by most cellular cargoes [8–10]. The karyopherins mediate active nuclear accumulation of the STAT dimers, leading to antiviral IFN-stimulated gene (ISG) activation [11]. To evade IFN-dependent immune signalling, viruses encode IFN-antagonist proteins, many of which target STAT1/STAT2, including through interactions leading to sequestration, induction of degradation and inhibition of phosphorylation [5]. Among IFN-antagonists, EBOV VP24 uses an unusual mechanism of competitive binding at the non-classical STAT1-binding site in NPI-1 karyopherins, thereby preventing STAT1 nuclear trafficking and ISG induction [12–15].

While IFN-STAT1/2 antagonism is reasonably well understood for many viruses, antagonism of other STATs including STAT3, the major mediator of signalling by IL-6 family cytokines (e.g. IL-6, oncostatin-M (OSM) [11]), is poorly defined, with only four mononegaviruses (three paramyxoviruses and one rhabdovirus) shown to express IFN-antagonist proteins that interact with STAT3 [16–19]. Nevertheless, STAT3-regulated processes are strongly implicated/dysregulated in EBOV disease, including the pro-inflammatory response, coagulation pathway and wound healing [6, 20–22]. Notably, despite critical roles in processes such as growth, development, apoptosis, infection and cancer, the precise mechanism(s) underlying cytokine-dependent STAT3 nuclear accumulation also remain poorly understood. Contrasting reports suggest three models whereby: (i) STAT3 undergoes constitutive nucleocytoplasmic shuttling with cytokines inducing intra-nuclear sequestration [23, 24], (ii) cytokine activation induces interaction of STAT3 with karyopherins including Kα1 resulting in nuclear import similar to STAT1 [25, 26], and (iii) STAT3 uses a combination of these mechanisms [27]. Notably, pY-STAT3 forms homodimers as well as heterodimers with pY-STAT1, which may regulate distinct gene subsets [28] and could use different trafficking mechanisms, possibly accounting for the contrasting models; this has not been directly examined.

Here, we aimed to examine the effect of EBOV on STAT3 responses, showing for the first time that EBOV VP24 antagonises STAT3 using a combination of mechanisms analogous to and distinct from that used for STAT1, to inhibit both STAT3 homodimers and heterodimers. We further reveal that the STAT3 complexes use distinct mechanisms for nuclear accumulation, apparently necessitating VP24’s multipronged strategy.

## Results and Discussion

### EBOV VP24 inhibits STAT3 responses

Despite likely roles in EBOV infection for dysregulation of cytokines/STATs other than IFN/STAT1/2, antagonism of other STATs by EBOV remains unresolved. To determine whether EBOV affects STAT3, we infected COS7 cells with EBOV before treatment with OSM [18, 25] and analysis of STAT3 localisation by immunofluorescence staining and confocal laser scanning microscopy (CLSM; Figure 1A). In mock-infected cells, STAT3 was diffusely localised between the nucleus and cytoplasm of resting cells, with nuclear accumulation clearly observed following OSM treatment, as expected. In EBOV-infected cells, however, OSM-dependent STAT3 nuclear accumulation was inhibited, with quantitative image analysis confirming a significant decrease in nucleocytoplasmic localisation in EBOV-compared with mock-infected cells (Figure 1A,B). To exclude possible impact by virus-induced type-I IFN, we confirmed that EBOV also antagonises STAT3 responses in Vero cells, which do not produce IFN (Figure 1A,B). Notably, in infected cells, STAT3 accumulated into distinct cytoplasmic regions (zoom images, Figure 1A), which EBOV nucleoprotein immunolabelling indicated to be viral inclusion/replication bodies (Figure S1). The finding of STAT3 accumulation into cytoplasmic viral inclusions is, to our knowledge, the first such observation for any virus.

**Figure 1.**
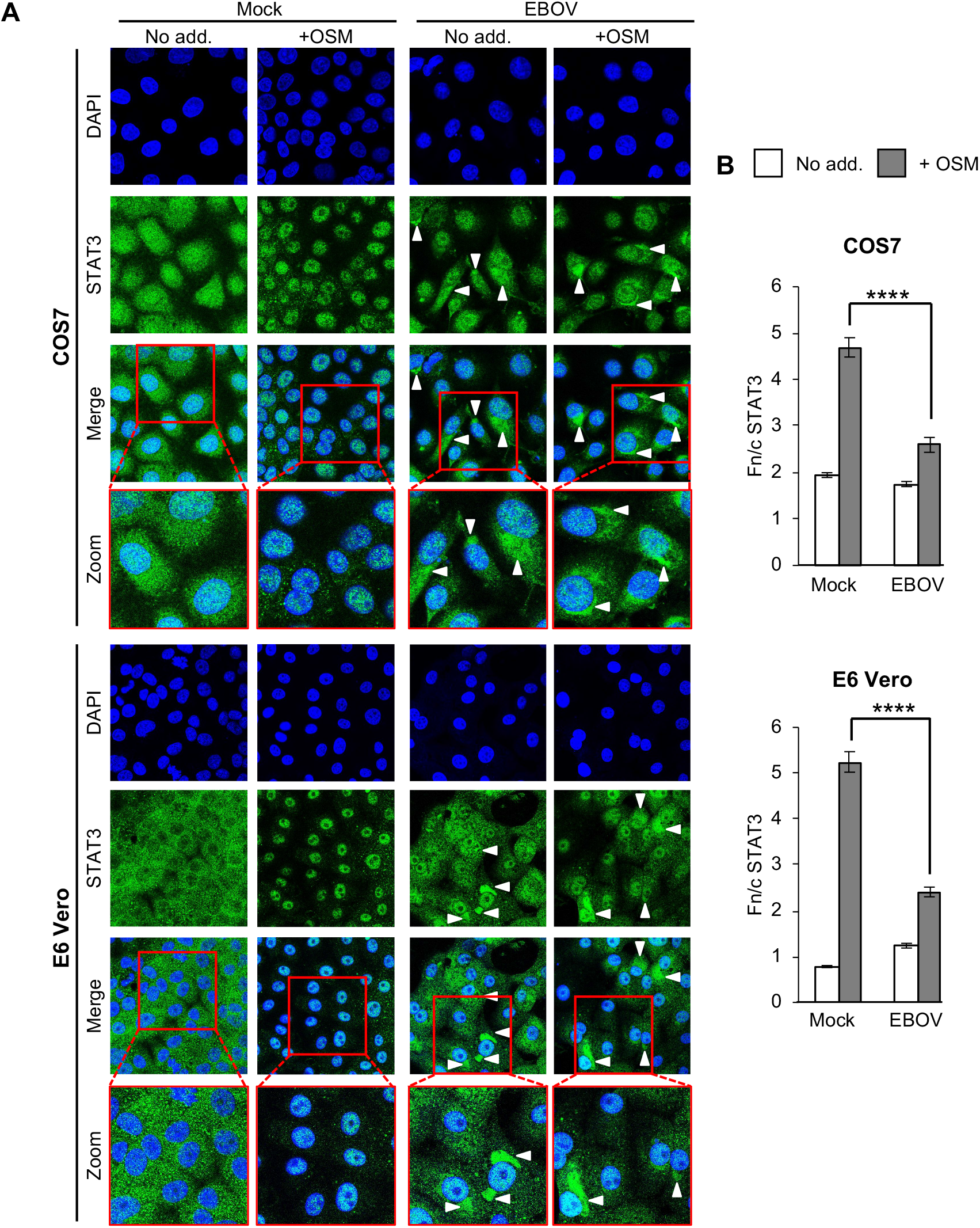
EBOV infection inhibits STAT3 responses to OSM. (A) COS7 (upper panel) or E6 Vero (lower panel) cells infected with EBOV (MOI 10, which results in infection of c. 100 % of cells, see Figure S1) or mock-infected were treated 72 h post-infection with or without OSM (10 ng/ml, 15 min) before fixation, immunofluorescent staining for STAT3 (green), and analysis by CLSM. DAPI (blue) was used to localise nuclei. Representative images are shown. Arrowheads indicate accumulation of STAT3 in cytoplasmic regions corresponding to viral inclusions (see Figure S1); indicated regions in merged images are expanded in panels below (Zoom). (B) Images such as those shown in A were analysed to calculate the nuclear to cytoplasmic fluorescence ratio (Fn/c) for STAT3 (mean ± SEM, n ≥ 70 cells for each condition). Statistical analysis (Student’s *t*-test) was performed using GraphPad Prism software; ****, p < 0.0001; No add., no addition.

Since VP24 antagonises IFN/STAT1 responses [12], we tested its effects on STAT3 by analysing COS7 cells expressing GFP-VP24 or negative controls (GFP or GFP-rabies virus (RABV) N-protein, which does not affect STAT3 [18]), and co-transfected to express mCherry-STAT3 (for live-cell analysis; Figure 2) or immunostained for endogenous STAT3 (Figure 3A,B). OSM effected clear nuclear accumulation of STAT3 in GFP and N-protein-expressing cells, but this was strongly inhibited in VP24-expressing cells. Since OSM can induce pY-STAT3 homodimers and pY-STAT3-pY-STAT1 heterodimers [29] and VP24 antagonises pY-STAT1 [12], we assessed the dependence of VP24-STAT3 antagonism on STAT1 using STAT1-deficient U3A cells [30, 31]. VP24 clearly antagonised STAT3 in U3A cells (Figure 3A,B) in which we confirmed a lack of STAT1 expression (Figure 3C), indicating that VP24 can inhibit STAT3 independently of STAT1 and thus antagonise STAT3 homodimers.

**Figure 2.**
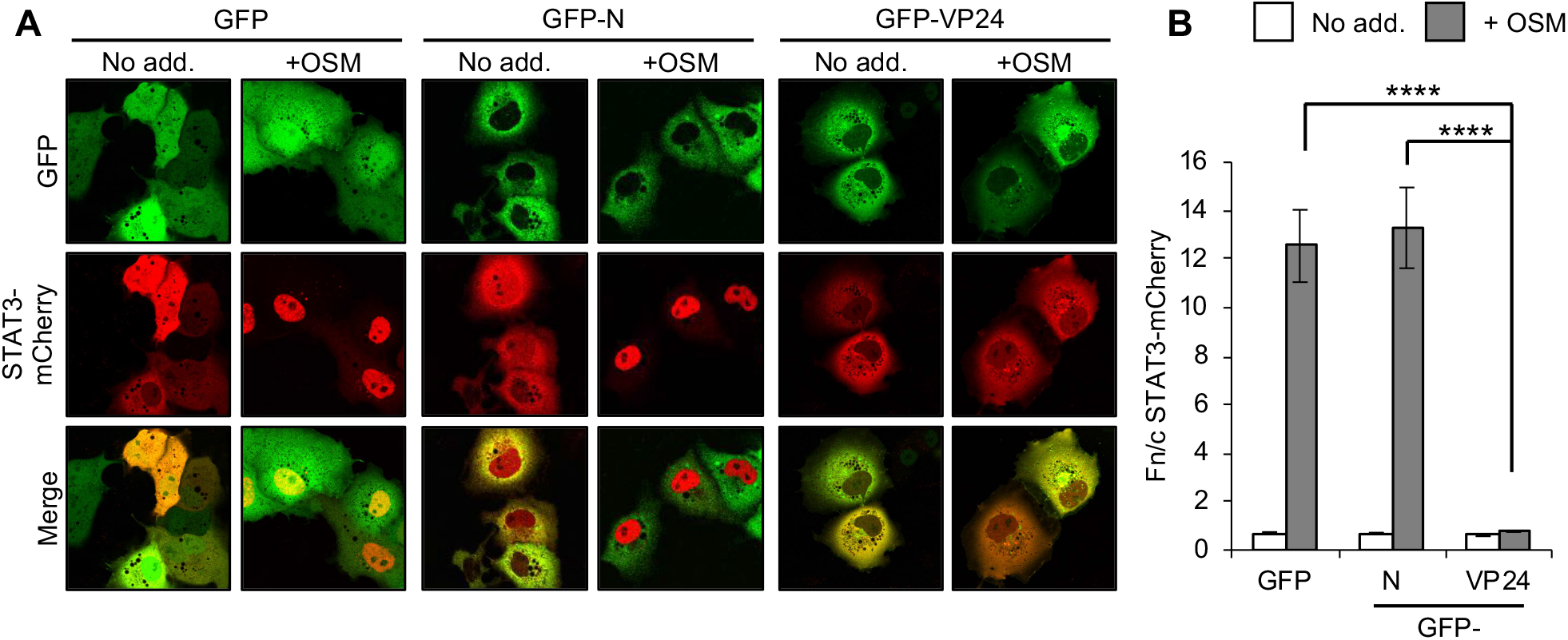
EBOV VP24 protein expression inhibits STAT3 responses to OSM. COS7 cells co-transfected to express the indicated proteins were treated 24 h post-transfection with or without OSM (10 ng/ml, 30 min) before live-cell CLSM analysis (A) to determine the Fn/c for STAT3-mCherry (B; mean ± SEM; n ≥ 35 cells for each condition; results are from a single assay representative of two independent assays; GFP-N, GFP-RABV-N-protein). Statistical analysis used Student’s *t*-test; ****, p < 0.0001.

**Figure 3.**
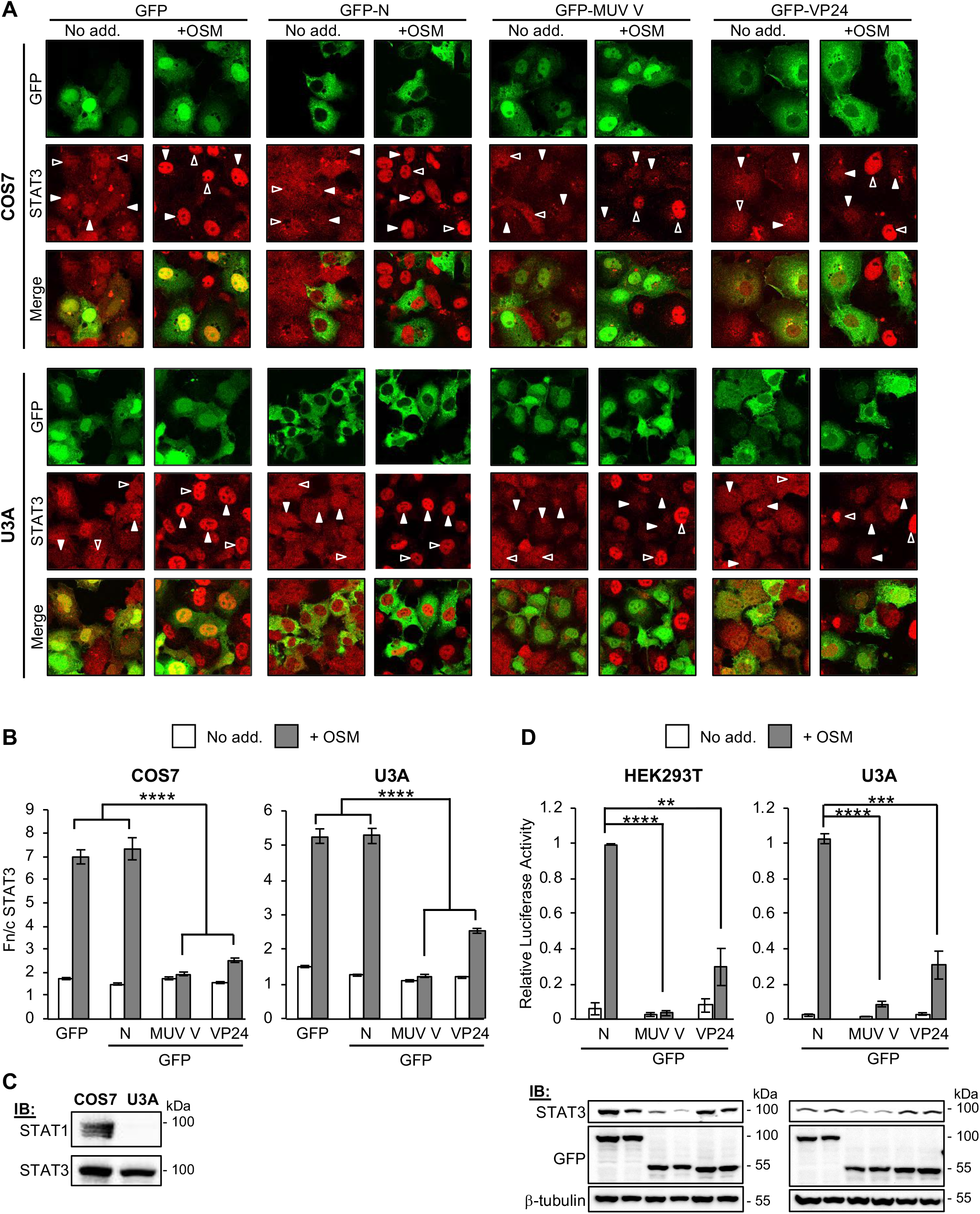
EBOV VP24 antagonises STAT3 independently of STAT1. (A,B) COS7 (upper panel) or U3A (lower panel) cells transfected to express the indicated proteins were treated 24 h post-transfection with or without OSM (10 ng/ml, 15 min) before fixation, immunofluorescent staining for STAT3 (red) and CLSM (A) to determine the Fn/c for STAT3 (B; mean ± SEM, n ≥ 35 cells for each condition; results are from a single assay representative of two independent assays). Filled and unfilled arrowheads indicate cells with or without, respectively, detectable expression of the transfected protein. MUV V, Mumps virus V protein. (C) Lysates of COS7 and U3A cells were analysed by immunoblotting (IB) for STAT1 and STAT3. (D) *upper panel*: HEK293T or U3A cells co-transfected with m67-LUC and pRL-TK plasmids, and plasmids to express the indicated proteins, were treated 16 h post-transfection with or without OSM (10 ng/ml, 8 h) before determination of relative luciferase activity (mean ± SEM; n = 3 independent assays); *lower panel:* cell lysates used in a representative assay were analysed by IB using antibodies against the indicated proteins. Statistical analysis used Student’s *t*-test; **, p<0.01; ***, p < 0.001; ****, p < 0.0001.

Analysis of OSM-dependent signalling using a luciferase reporter gene assay [18, 32] in HEK293T and U3A cells indicated that VP24 effects significant suppression of OSM/STAT3 signalling (Figure 3D; upper panel); RT-qPCR analysis confirmed that VP24 can inhibit OSM-induced expression of the STAT3-dependent *socs3* gene (Figure S2). Mumps virus V-protein (MUV-V, used as a positive control in our assays) induces STAT3 degradation to suppress IL-6 signalling [16]. We confirmed that MUV-V inhibits STAT3 responses and that this correlates with reduced levels of STAT3 expression in cell lysates. Since no similar effect was observed on STAT3 expression in VP24-expressing cells (Figure 3D; lower panel), it appeared that VP24 uses a different antagonistic mechanism.

### VP24 inhibits Kα1 interaction with STAT3, dependent on STAT1

VP24 antagonises STAT1 responses by competitive binding to karyopherins [12, 13, 15], including Kα1 that is also reported to mediate STAT3 nuclear import [25–27]. We thus examined whether VP24 can displace STAT3 from Kα1, by immunoprecipitation of FLAG-Kα1 from OSM-treated HEK293T cells (as previously used to analyse effects of VP24 on IFN-activated pY-STAT1-karyopherin interactions [12, 13]) or U3A cells. Cells were co-transfected to express FLAG-Kα1 with GFP-VP24 or GFP, before OSM treatment and lysis for IP (Figure 4). pY-STAT1, pY-STAT3 and GFP-VP24 co-precipitated specifically with Kα1 as expected, consistent with reports that STAT1 and STAT3 are Kα1 cargoes [8, 25, 26], and VP24 can interact with Kα1 [12, 13]. Clearly, for both pY-STAT1 (as expected [12, 13]) and pY-STAT3, the amount co-precipitated with Kα1 from HEK293T cells was reduced by VP24, consistent with competitive binding. Importantly, although a number of IFN-antagonists suppress STAT phosphorylation [5], VP24 did not affect levels of pY-STAT1 or pY-STAT3 in lysates, indicating that reduced Kα1 interaction of STAT3 is not due to altered phosphorylation. Thus, it appears that VP24 can compete with STAT3-containing complexes for Kα1 interaction, similarly to its effect on STAT1.

**Figure 4.**
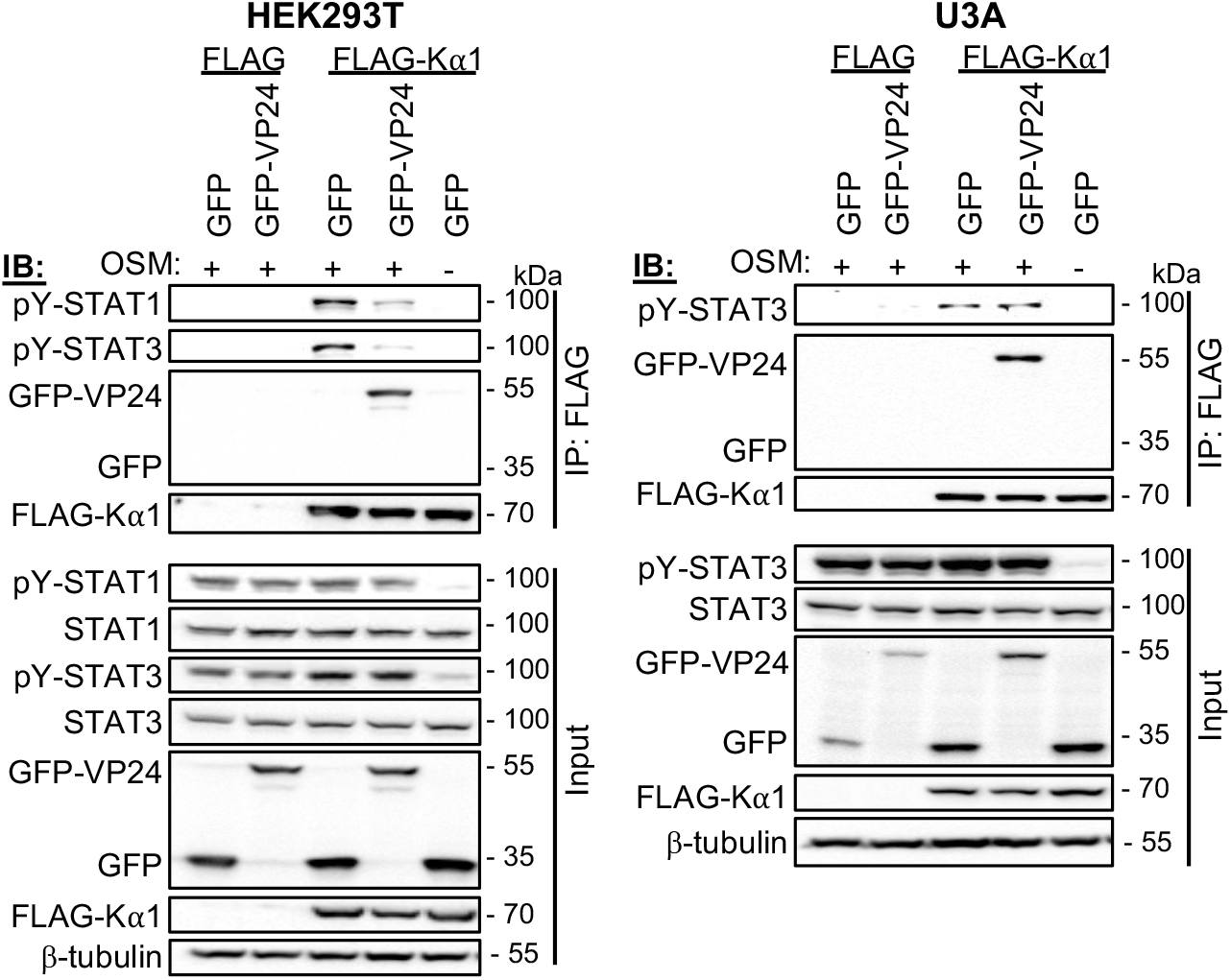
EBOV VP24 inhibits Kα1-STAT3 interaction, dependent on STAT1. HEK293T or U3A cells co-transfected to express the indicated proteins were treated 24 h post-transfection with or without OSM (10 ng/ml, 15 min) before lysis and immunoprecipitation for FLAG. Lysates (input) and immunoprecipitates (IP) were analysed by IB using antibodies against the indicated proteins. Results are representative of ≥ 2 independent assays. Expanded images of all membranes are shown in Figure S4.

Intriguingly, however, co-immunoprecipitation assays in U3A cells indicated that VP24 does not affect Kα1-pY-STAT3 interaction (Figure 4), despite clear impact on STAT3 responses in these cells (Figure 3). It has been suggested that karyopherin interactions of STAT homo- and heterodimers might differ [23, 24], and our data support this, providing evidence that STAT3 homodimers may form interactions at a site in the karyopherin distinct to the non-classical STAT1/VP24-binding site, while STAT3-STAT1 heterodimers appear to bind at the STAT1/VP24 site and so can be displaced by VP24 (Figure 4). Since STAT1 homodimers and STAT1-STAT2 heterodimers also bind to this site [10], this might represent a common interface for STAT1-containing complexes, such that competitive binding by VP24 is likely to occur for heterodimers activated by other cytokines/mediators (e.g. STAT4-STAT1 heterodimers). Since the data from U3A cells indicate that STAT3 homodimers bind to Kα1 *via* a site not bound by VP24, it appears that an alternative mechanism must antagonise signalling by these complexes.

### VP24 does not inhibit STAT3 binding to DNA

Reports supporting constitutive nuclear trafficking of STAT3 suggest that STAT3 accumulates in the nucleus in response to cytokine due to intra-nuclear interactions/sequestration, such as through induced DNA binding [24]. We therefore considered that VP24 may inhibit STAT3 nuclear accumulation in U3A cells by inhibiting the capacity of STAT3 to bind DNA, similar to the antagonistic mechanism of RABV P-protein for STAT1, where the P-protein binds proximal to or within the STAT1 DNA binding domain [33, 34]. To assess DNA binding by STAT3 directly, we performed electrophoretic mobility shift assay (EMSA) analysis of cell lysates using the m67 probe (Figure 5), which is a high affinity variant of the sis-inducible element from the *C-FOS* gene, commonly used to analyse STAT3-DNA binding [35–37]. OSM induced clear DNA binding of both endogenous and overexpressed STAT3 in the absence and presence of VP24, with VP24 having no evident inhibitory effect. Thus, the principal mechanism of antagonism does not appear to involve a direct hindrance of STAT3-DNA interaction.

**Figure 5.**
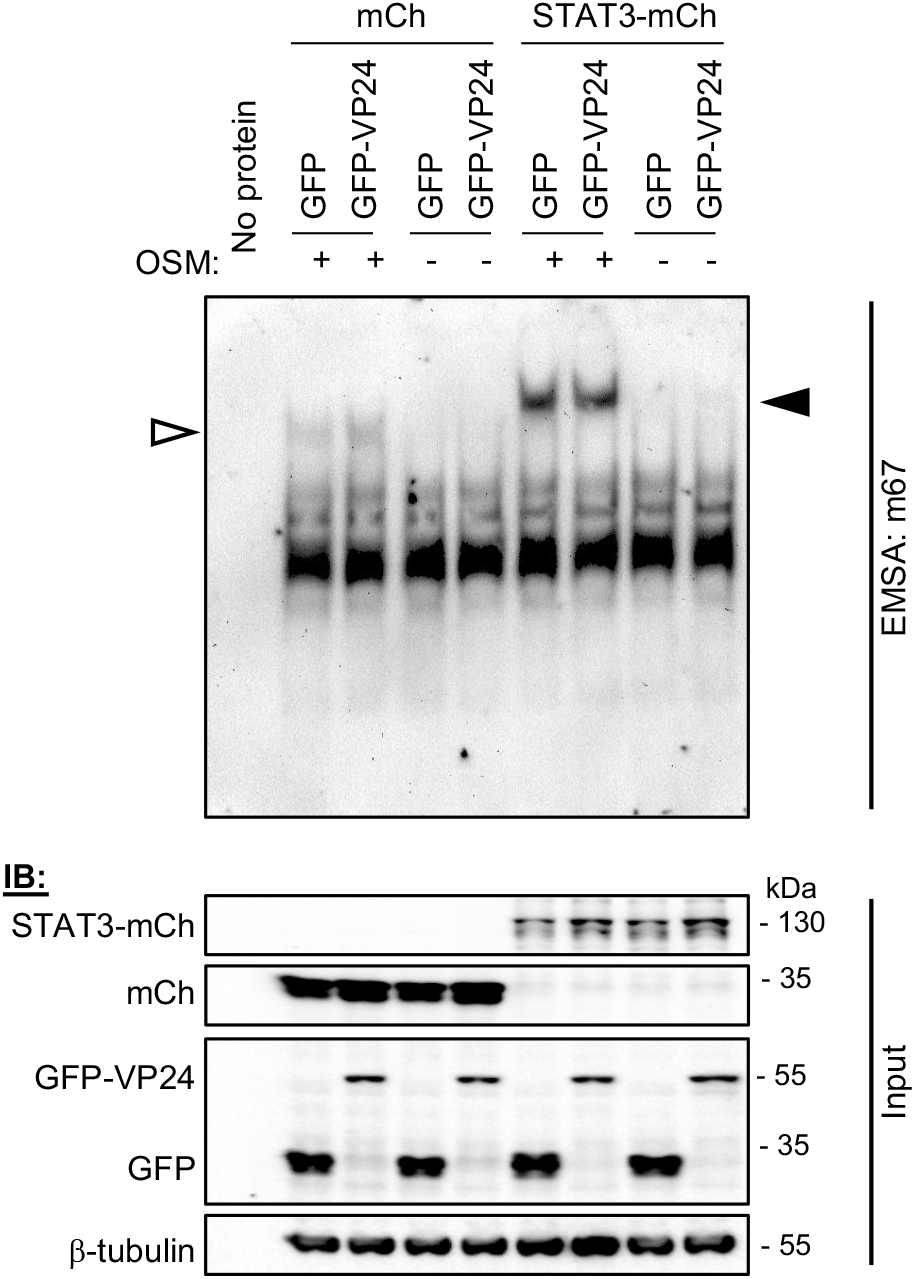
EBOV VP24 does not prevent interaction of STAT3 with target DNA. *Upper panel:* U3A cells co-transfected to express the indicated proteins were treated 24 h post-transfection with or without OSM (10 ng/ml, 15 min) before lysis and incubation of equal amounts of cell lysate protein or no lysate control (no protein) with digoxigenin-labelled m67 probe. EMSA reactions were resolved on 4.5 % polyacrylamide gel in 0.5 x TBE, before transfer to a nylon membrane and IB for digoxigenin. Results are representative of 3 independent assays. Filled and unfilled arrowheads indicate bands consistent with DNA complexes with STAT3-mCherry and endogenous STAT3, respectively. *Lower panel:* Cell lysates were also analysed by SDS-PAGE and IB (input) using antibodies against the indicated proteins.

### VP24 interacts with STAT3, independently of VP24-karyopherin binding

Recombinant purified VP24 and STAT1 proteins were reported to interact *in vitro* [38], but no direct interaction has been detected for proteins expressed in mammalian cells, so there is currently no evidence that this is significant to STAT1 antagonist function [15, 39]. Nevertheless, since STAT3 localizes into viral inclusion bodies (Figure 1A), of which VP24 is a component [40], and many IFN-antagonists inhibit STATs through physical interaction [5], we tested whether VP24 can bind to STAT3. Endogenous and transfected STAT3 co-precipitated with VP24 from U3A cells (Figure 6A,B), and reciprocal immunoprecipitation *via* STAT3 confirmed the interaction (Figure S3). Thus, VP24 interacts with STAT3 independently of STAT1, consistent with data for antagonism of OSM/STAT3 signalling (Figure 3). We also confirmed co-precipitation of STAT3 with VP24 from HEK293T and COS7 cells (Figure 6C, Figure S6).

**Figure 6.**
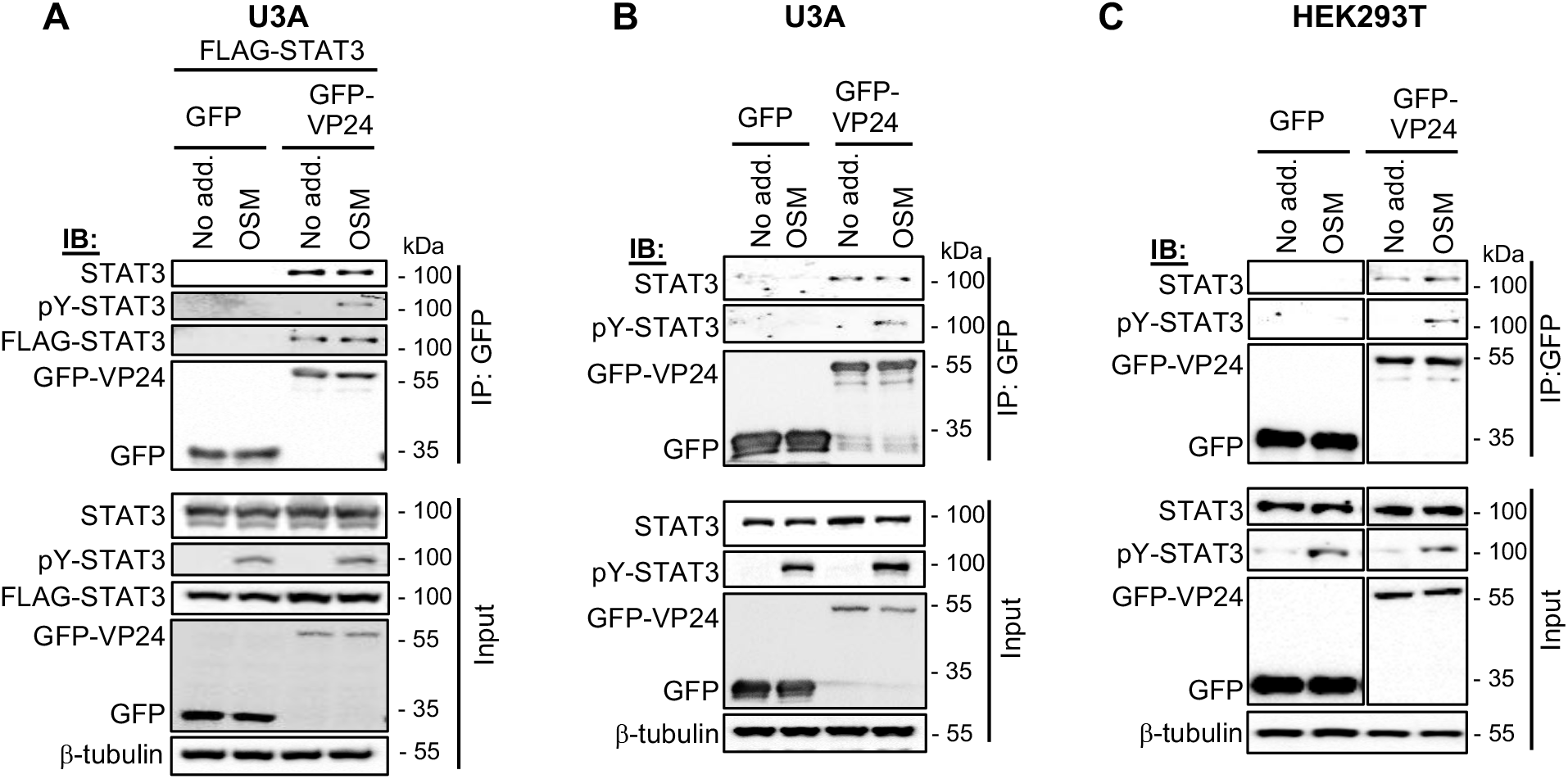
EBOV VP24 interacts with STAT3. (A) U3A cells co-transfected to express FLAG-STAT3 and GFP or GFP-VP24 as indicated were treated 24 h post-transfection with or without OSM (10 ng/ml, 30 min) before lysis, immunoprecipitation for GFP, and IB, as described in the legend to Figure 4. (B,C) U3A (B) or HEK293T (C) transfected to express the indicated proteins were treated with or without OSM (10 ng/ml, 15 min) before immunoprecipitation for GFP and IB for endogenous STAT3. Results are representative of 2 independent assays and show data from a single blot with intervening and marker lanes removed. Expanded images of all membranes are shown in Figures S5-S6.

To further investigate the antagonistic mechanism, we analysed a karyopherin-binding deficient VP24 protein, wherein mutations of key residues at the VP24:karyopherin interface (MUT; L201A/E203A/P204A/D205A/S207A) strongly impair karyopherin binding and STAT1/IFN antagonism [15]. The effect of the mutations in inhibiting STAT1-antagonist function was confirmed using a STAT1/2-IFN-dependent luciferase reporter assay (using pISRE-LUC plasmid), which indicated an almost nine fold increase in luciferase activity in IFN-α-treated HEK293T cells expressing mutated protein compared with wild-type (WT) protein (Figure 7A; left panel). Interestingly, analysis using the STAT3/OSM-dependent signalling assay (using m67-LUC plasmid) in U3A cells showed no significant impact of the mutations on VP24 inhibitory activity (Figure 7A; middle panel), indicating that specific antagonism of STAT3 by VP24 is independent of karyopherin-binding activity, consistent with the lack of an effect of VP24 on Kα1-STAT3 interaction in U3A cells. Assays of STAT3/OSM-dependent signalling in HEK293T cells, however, indicated some dependence on karyopherin-binding, probably reflecting a contribution to signalling by STAT3-STAT1 heterodimers (Figure 7A; right panel), and consistent with the capacity of VP24 to compete with STAT1 and STAT3 for Kα1 interaction in these cells. Thus, it appears that in contrast to STAT1 (and STAT3-STAT1 heterodimers), antagonism of signalling by STAT3 homodimers is independent of VP24-karyopherin binding. Consistent with this, the mutations had no evident effect on VP24-STAT3 interaction in U3A cells (Figure 7B). Taken together, these data indicate that the nuclear trafficking mechanisms of STAT1 and STAT3 are distinct, and, accordingly, antagonism by VP24 uses different mechanisms, likely including competition with STAT1-containing complexes for karyopherin binding, as well as physical interaction with STAT3, which can cause localisation into cytoplasmic inclusions.

**Figure 7.**
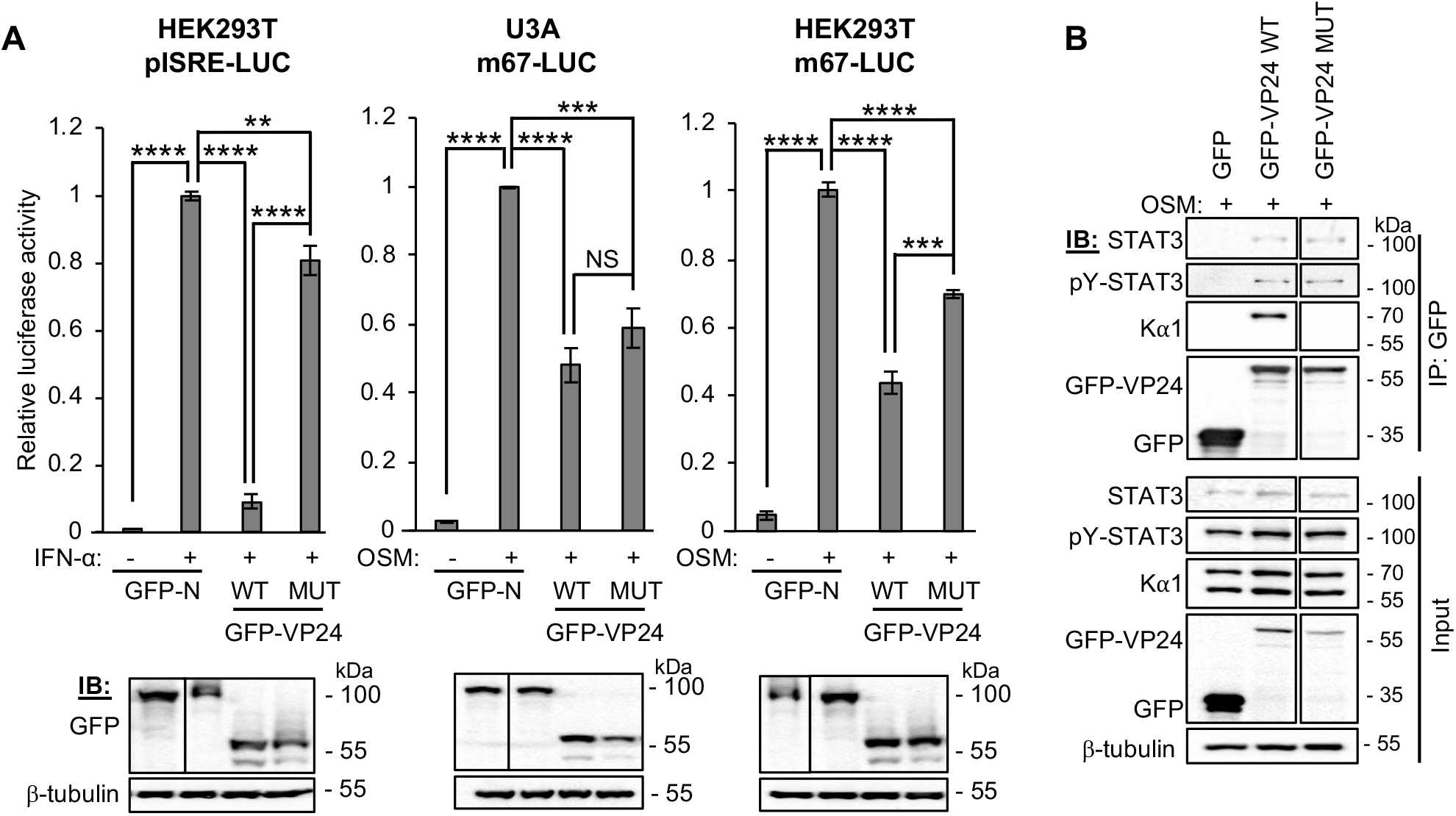
Antagonism of STAT3 by EBOV VP24 in U3A cells is independent of VP24-karyopherin interaction. (A) *upper panel:* HEK293T or U3A cells co-transfected with pISRE-LUC or m67-LUC plasmid, pRL-TK plasmid, and plasmids to express the indicated proteins, were treated 8 h (IFN-α) or 16 h (OSM) post-transfection with or without IFN-α (1,000 U/ml for 16 hours) or OSM (10 ng/ml for 8 h) before determination of relative luciferase activity (mean ± SEM; n ≥ 3 independent assays); *lower panel:* cell lysates used in representative assays were analysed by IB for GFP and β-tubulin. Statistical analysis used Student’s *t*-test; **, p < 0.01; ***, p < 0.001; ****, p < 0.0001; NS, not significant. (B) U3A cells transfected to express the indicated proteins were treated with OSM before immunoprecipitation for GFP and IB, as described in the legend to Figure 6. Results are representative of 2 independent assays and show data from a single blot with intervening and marker lanes removed. Expanded images of membranes are shown in Figure S7.

These findings indicate that VP24 uniquely uses two distinct mechanisms to inhibit different STAT3 complexes, consistent with important roles for global shutdown of STAT3 in EBOV infection, possibly relating to the dysregulation of inflammation, coagulation and mucosal wound healing observed during EBOV infection [6, 20–22, 41, 42]. Recent reports indicate that STAT3 antagonism by MUV is associated with neurovirulence *in vivo* [43], and suppression of IL-6 signalling by influenza A virus early in infection contributes to a cytokine storm implicated in disease severity [44]. Interestingly, although the IFN-antagonist VP40 of the filovirus Marburg virus does not specifically target STATs, it inhibits upstream kinases resulting in inhibition of activation of both STAT1 and STAT3 [45]. Together these data indicate that potent suppression of STAT3 responses by filoviruses may contribute to excessive inflammatory responses associated with severe haemorrhagic fever. The apparent importance of STAT3 targeting to filoviruses, and previous reports of roles in infection by paramyxoviruses and rhabdoviruses, also indicates that specific and direct antagonism of STAT3 is important to diverse pathogens in the order *Mononegavirales* [16–19]. Taken together, these data suggest that virus-STAT3 interactions could provide potential targets for antivirals for diverse pathogens. Beyond the implications for viral infection, the study also provides, to our knowledge, the first clear indication of distinct nuclear import strategies for STAT3 homodimers and heterodimers. This potentially accounts for the contrasting trafficking models previously proposed [23–27], and supports the idea that these complexes have distinct roles in signalling by STAT3, a pleiotropic molecule important to processes including cancer, development and immunity.

## Materials and Methods

### Plasmids and Cell Culture

Constructs to express EBOV-VP24 and MUV-V fused to GFP were generated by PCR amplification from pCAGGS-FLAG-VP24 (kindly provided by Christopher Basler, Georgia State University) and MUV V-FLAG (a gift from Curt Horvath [16], Addgene plasmid #44908), and cloning into the pEGFP-C1 vector C-terminal to GFP (Clontech). Constructs to express mCherry- or FLAG-tagged STAT3 were kind gifts from Marie Bogoyevitch (University of Melbourne), and the construct to express FLAG-tagged Kα1 was a kind gift from Christopher Basler (Georgia State University). Other constructs have been described elsewhere [18, 32]. U3A (a kind gift from George Stark, Lerner Research Institute, Cleveland Clinic), COS7, E6 Vero and HEK293T cells were maintained in DMEM supplemented with 10 % FCS and GlutaMAX (Life Technologies), 5 % CO2, 37oC. Transfections used Lipofectamine 2000 (Invitrogen), Lipofectamine 3000 (Invitrogen), or FuGene HD (Promega), according to the manufacturer’s instructions.

### Virus infection

All infectious work was conducted at Physical Containment Level 4 (PC4) at the Australian Centre for Disease Preparedness (ACDP, formerly AAHL). EBOV infections used Mayinga 1976 isolate (MOI of 10), which was originally received from NIH Rocky Mountain Laboratories and passaged three times in Vero cells at ACDP after receipt.

### CLSM

For analysis of STAT3 localisation, cells growing on coverslips transfected with plasmids or infected with EBOV were incubated in serum-free-(SF)-DMEM for 1 h and treated without or with 10 ng/mL recombinant human OSM (BioVision) for 15 min (analysis of fixed/immunostained cells) or 30 min (analysis of STAT3-mCherry in living cells) before fixation using 3.7 % formaldehyde (10 min, room temperature (RT) for transfected cells) or 4 % paraformaldehyde (48 h, 4oC for infected cells), followed by 90 % methanol (5 min, RT) and immunostaining. Antibodies used for were: anti-STAT3 (Santa Cruz, sc-482; or Cell Signaling Technology, 9139), anti-EBOV nucleoprotein (rabbit clone #691, final bleed 1410069), anti-mouse Alexa Fluor 488 (ThermoFisher Scientific, A11001) and anti-rabbit Alexa Fluor 568 (ThermoFisher Scientific, A11036). Imaging used a Leica SP5 or Nikon C1 inverted confocal microscope with 63 X objective. For live cell analysis, cells were imaged in phenol-free DMEM using a heated chamber. Digitized confocal images were processed using Fiji software (NIH). To quantify nucleocytoplasmic localisation, the ratio of nuclear to cytoplasmic fluorescence, corrected for background fluorescence (Fn/c), was calculated for individual cells expressing transfected protein [18, 32]; mean Fn/c was calculated for n ? 35 cells for each condition in each assay.

### Co-immunoprecipitation

Transfected cells were incubated in SF-DMEM (3 h) before treatment with or without OSM (10 ng/ml, 15 min), lysis and immunoprecipitation using GFP-Trap Agarose beads (Chromotek) or Anti-FLAG M2 Magnetic beads (Sigma-Aldrich), according to the manufacturer’s instructions. Lysis and wash buffers were supplemented with PhosSTOP (Roche), cOmplete Protease Inhibitor Cocktail (Roche) and 10 mM NaF. Lysates and immunoprecipitates were analysed by SDS-PAGE and immunoblotting (IB) using antibodies against STAT3 (above), pY-STAT3 (Cell Signaling Technology, 9145), STAT1 (Cell Signaling Technology, 14994), pY-STAT1 (Tyr701, Cell Signaling Technology, 9167), FLAG (Sigma-Aldrich, F1804), GFP (Roche Applied Science, 11814460001), mCherry (Abcam, ab167453), Kα1 (Abcam, ab154399) and β-tubulin (Sigma-Aldrich, T8328), and HRP-conjugated secondary antibodies (Merck). Visualisation of bands used Western Lightning chemiluminescence reagents (PerkinElmer).

### Luciferase Reporter Gene Assays

Cells were co-transfected with m67-LUC or pISRE-LUC (in which Firefly luciferase expression is under the control of a STAT3 or STAT1/2-dependent promoter, respectively) and pRL-TK (transfection control, from which *Renilla* luciferase is constitutively expressed), as previously described [18, 46], together with protein expression constructs. Cells were treated 16 h (OSM) or 8 h (IFN-α) post-transfection with or without OSM (10 ng/mL for 8 h) or IFN-α (1,000 U/ml for 16 hours) before lysis using Passive Lysis Buffer (Promega). Firefly and *Renilla* luciferase activity was then determined in a dual luciferase assay, as previously described [18, 46]. The ratio of Firefly to *Renilla* luciferase activity was determined for each condition, and then calculated relative to that determined for GFP-N-protein-expressing cells treated with OSM (relative luciferase activity). Data from ≥ 3 independent assays were combined, where each assay result is the mean of three replicate samples.

### EMSA

Transfected cells were incubated in SF-DMEM (2 h) before treatment with or without OSM (10 ng/ml, 15 min) and lysis in 20 mM Hepes (pH 7.0), 300 mM NaCl, 20 % (v/v) glycerol, 10 mM KCl, 1 mM MgCl2, 0.5 mM DTT, 0.1 % (v/v) Triton X-100, as previously [47], supplemented with PhosSTOP (Roche), cOmplete Protease Inhibitor Cocktail (Roche) and 10 mM NaF. 10 ng of clarified cell lysate (calculated using Pierce Microplate BCA Protein Assay Kit - Reducing Agent Compatible, ThermoFisher Scientific) was incubated with 1 ng of digoxigenin-labelled m67 probe (double-stranded; 5’-AGCTTCATTTCCCGTAAATCCCTA-3’) in a reaction containing 20 mM Hepes (pH 7.6), 30 mM KCL, 10 mM (NH4)2SO4, 1 mM DTT, 1 mM EDTA, 0.2 % (w/v) Tween-20, 1 *μ*g poly[d(I-C)] and 0.1 *μ*g poly-Lysine (based on DIG Gel Shift Kit, 2nd Generation, Roche) for 15 min at RT. DNA-protein complexes were resolved on a 4.5 % polyacrylamide gel in 0.5 x TBE running buffer (4oC), before electrophoretic transfer to a nylon membrane and IB using anti-Digoxigenin-AP Fab fragments (Roche). Visualisation of bands used CDP-Star chemiluminescence reagents (Roche).

### RT-qPCR

Transfected HEK293T cells were incubated in SF-DMEM (3 h) before treatment without or with OSM (10 ng/ml, 45 min) and RNA extraction (ReliaPrep RNA Cell Miniprep System, Promega). cDNA was generated using oligo(dT)20 primer (GoScript Reverse Transcription System, Promega), before RT-qPCR using primers for *socs*3 and *gapdh*, and iTaq Universal SYBR Green Supermix (Bio-Rad). Standard curves were generated for each primer pair using serial dilutions of the reference cDNA (samples from GFP-N-protein-expressing cells treated with OSM). *Socs3* expression was normalized to *gapdh* [46], and then calculated relative to that for GFP-N-expressing cells treated with OSM. Data from 2 independent assays were combined, where each assay result is the mean of replicate samples. Primer sequences were: 5’-GGAGTTCCTGGACCAGTACG-3’ and 5’-TTCTTGTGCTTGTGCCATGT-3’ for *socs3;* 5’-GAAGGTGAAGGTCGGAGTC-3’ and 5’-GGTCATGAGTCCTTCCACGAT-3’ for *gapdh*.

### Statistical Analysis

Unpaired two-tailed Student’s *t*-test was performed using Prism software (version 7, GraphPad).

## Acknowledgements

We acknowledge Cassandra David for assistance with tissue culture, and the facilities and technical assistance of the Biological Optical Microscopy Platform (University of Melbourne) and Monash Micro Imaging Facility (Monash University). We also thank staff of the Pathology and Pathogenesis Group at ACDP (CSIRO) for assistance with microscopic examination of infected samples, and the Australian Microscopy and Microanalysis Research Facility for support with equipment within the ACDP microscopy facility. Plasmids to express FLAG-VP24 and FLAG-Kα1 were kind gifts from Christopher Basler (Georgia State University); plasmids to express STAT3 were kind gifts from Marie Bogoyevitch (University of Melbourne); plasmid to express Mumps V-FLAG was a kind gift from Curt Horvath (Addgene plasmid #44908). U3A cells were a kind gift from George Stark (Lerner Research Institute, Cleveland Clinic). We also thank Michelle Audsley for critical reading of the manuscript.

## Author Contributions

A.R.H. and G.W.M. designed experiments, analysed data and wrote the manuscript. A.R.H performed the experiments, except EBOV infection experiments, which were performed by S.T. and G.A.M, and preparation/imaging of infected samples, which was performed by M.D. and D.G.

