## Supplemental Figures for "Antagonism of STAT3 signalling by Ebola virus"

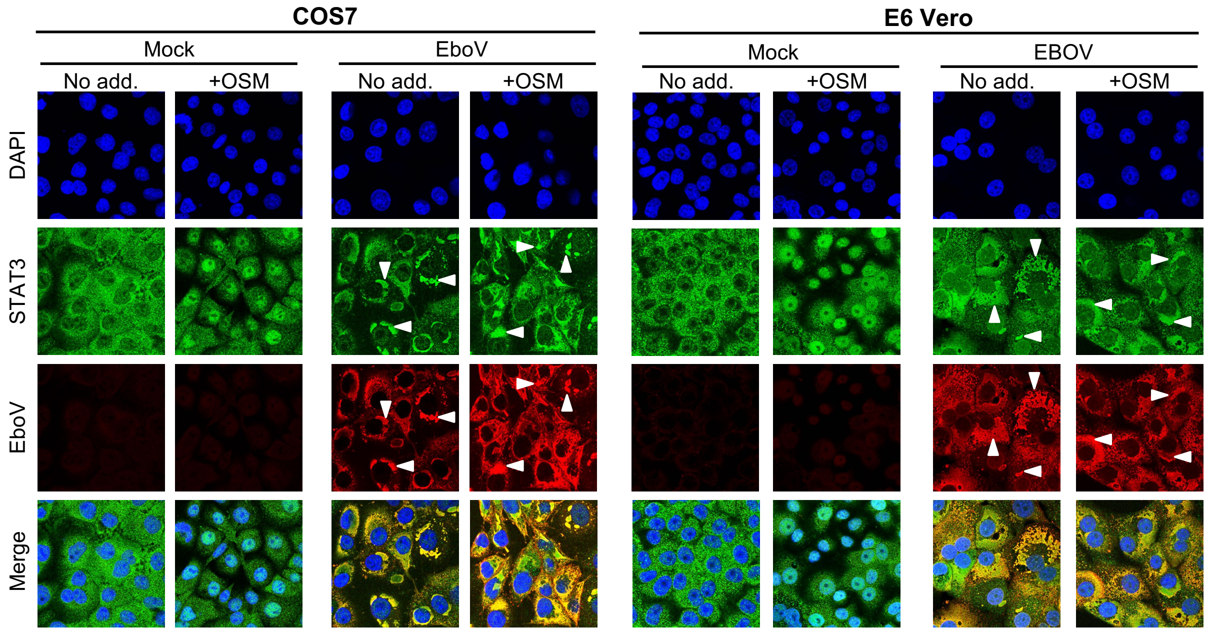


**Figure S1. Cytoplasmic regions of STAT3 accumulation in EBOV-infected cells correspond to EBOV protein-enriched inclusion bodies.** COS7 (left panel) or E6 Vero (right panel) cells infected with EBOV (MOI 10) were treated 72 h post-infection with or without OSM (10 ng/ml, 15 min) before fixation, immunofluorescent staining for EBOV nucleoprotein (red) and STAT3 (green), and analysis by CLSM. DAPI (blue) was used to localise nuclei. Images are representative of ≥ 5 fields of view for each condition. Arrowheads indicate accumulation of nucleoprotein and STAT3 in discrete cytoplasmic regions/inclusions.

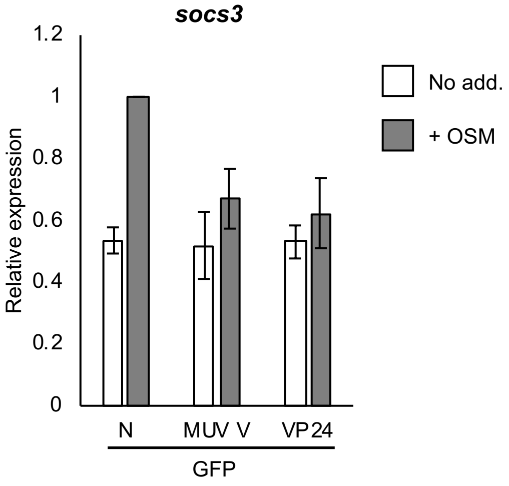


**Figure S2. VP24 inhibits STAT3-dependent gene expression.** HEK293T cells transfected to express the indicated proteins were treated 24 h post-transfection with or without OSM (10 ng/ml, 45 min) before analysis by RT-qPCR. Histogram shows expression of *socs3* calculated relative to *gapdh* and normalised to GFP-N-expressing cells treated with OSM (mean ± standard deviation; n = 2 independent assays).

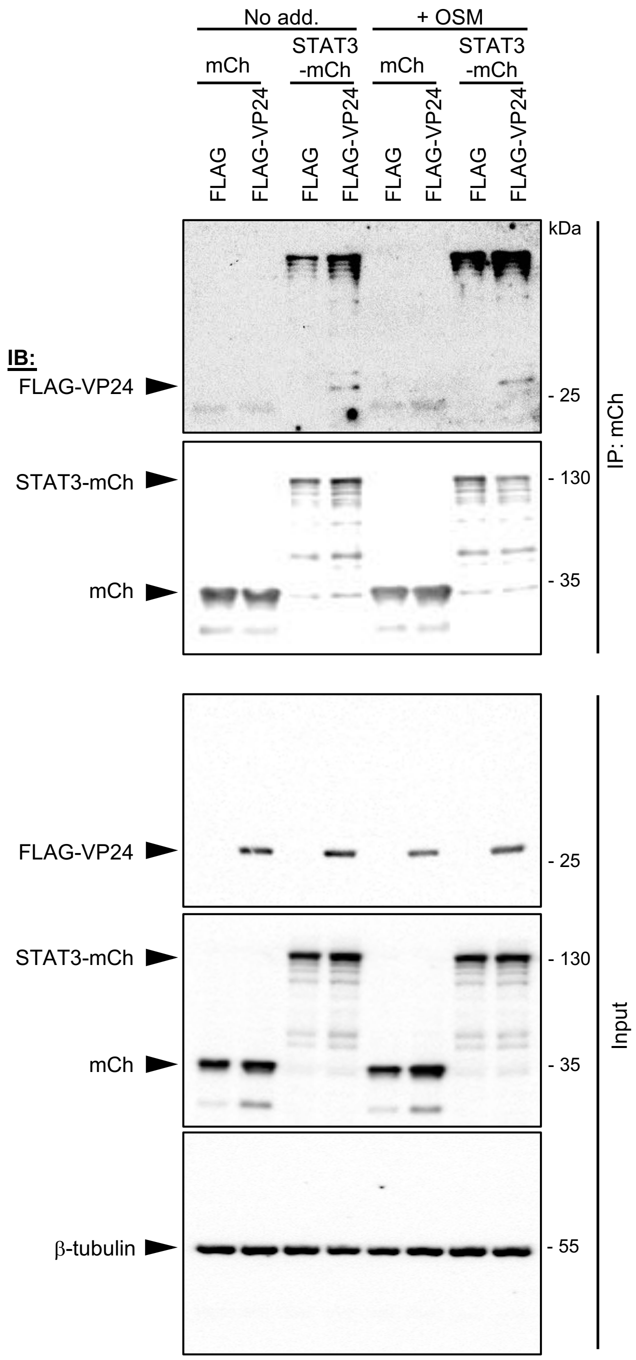


**Figure S3. Reciprocal immunoprecipitation of VP24 with STAT3.** U3A cells co-transfected to express STAT3-mCherry or mCherry and FLAG-VP24 or FLAG were treated with or without OSM before immunoprecipitation for mCherry and IB, as described in the legend to Figure 6A. Arrowheads indicate specific protein bands.

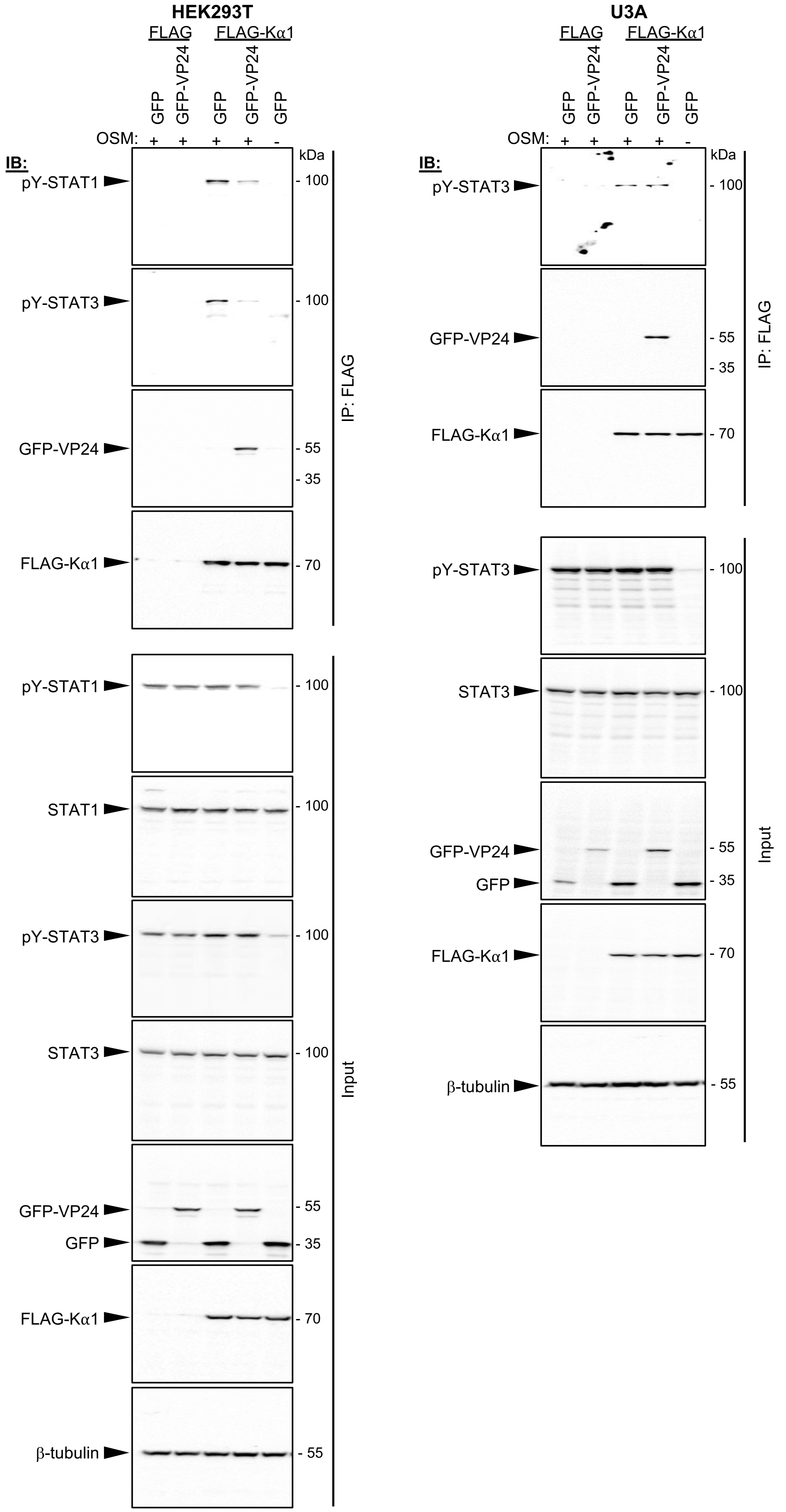


**Figure S4. Expanded images of western blots shown in Figure 4.** Full images of membranes shown in Figure 4; arrowheads indicate specific proteins bands.

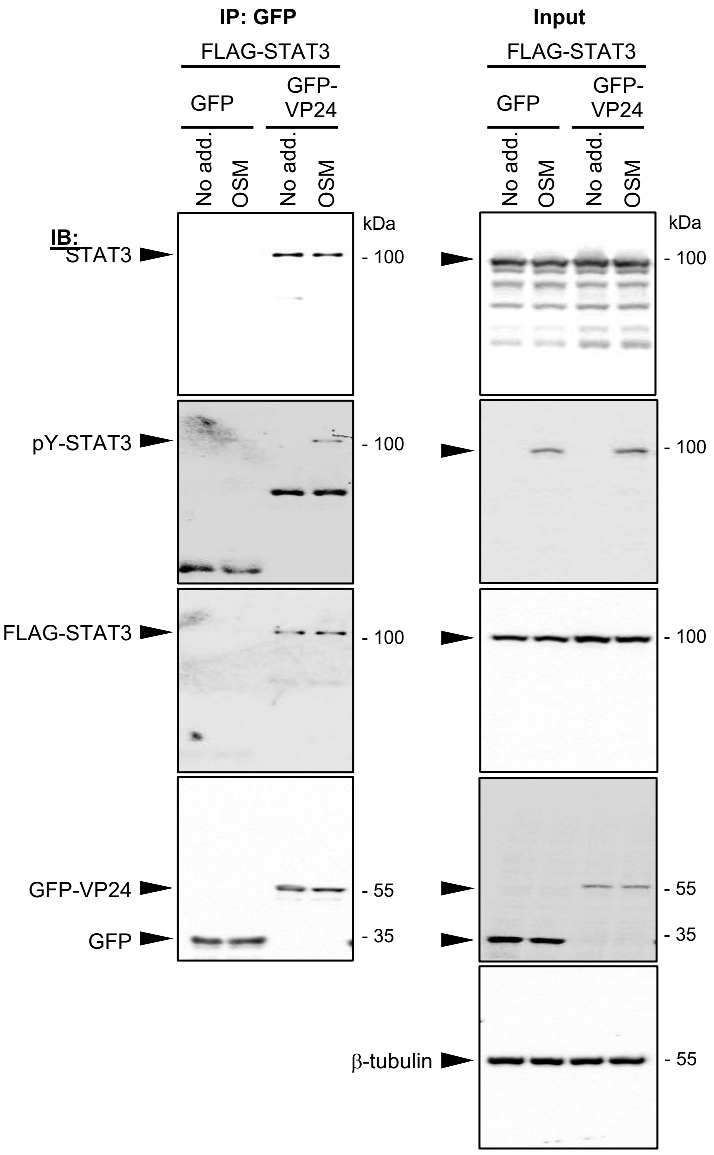


**Figure S5. Expanded images of western blots shown in Figure 6A.** Full images of membranes shown in Figure 6A; arrowheads indicate specific proteins bands.

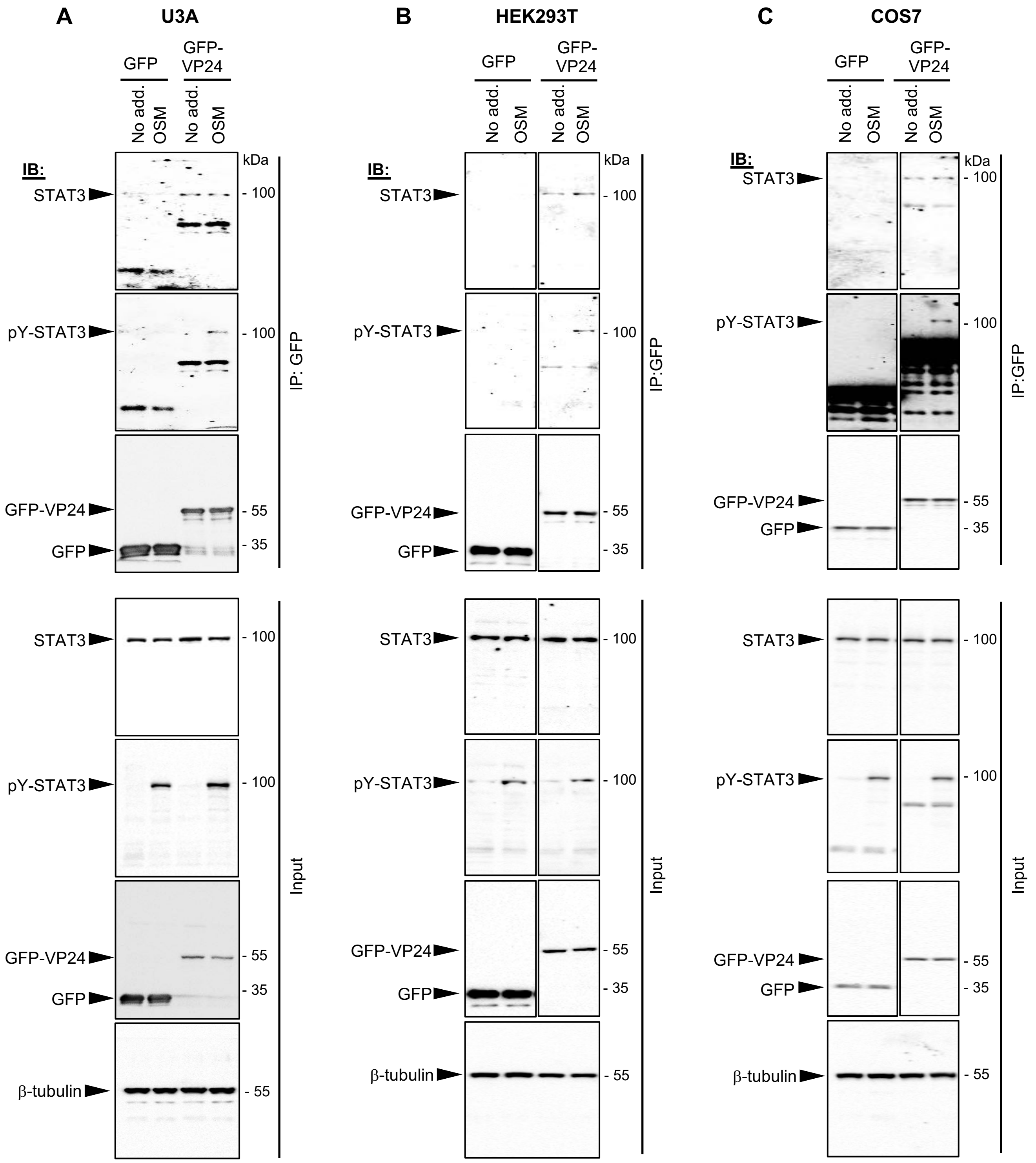


**Figure S6. Expanded images of western blots shown in Figure 6B-C and western analysis of immunoprecipitation assay using COS7 cells.** (A,B) Full images of membranes shown in Figure 6B (A) and Figure 6C (B). (C) Results of immunoprecipitation assay using COS7 cells transfected and treated as described for U3A and HEK293T cells in Figure 6B-C. Results are representative of 2 independent assays and show data from a single blot with intervening and marker lanes removed. Arrowheads indicate specific proteins bands.

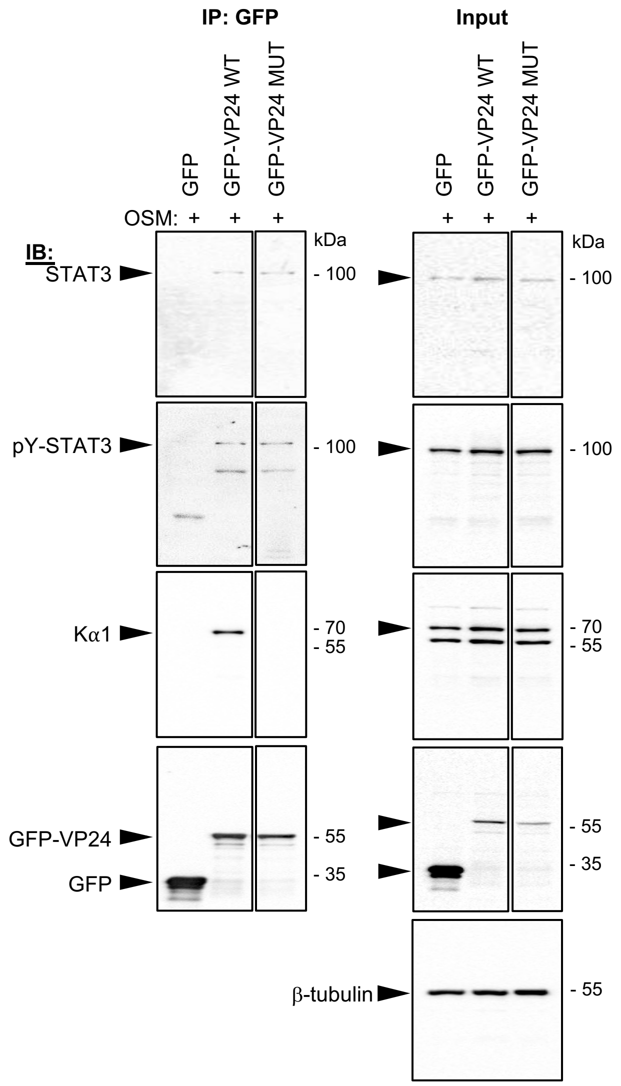


**Figure S7. Expanded images of western blots shown in Figure 7B.** Full images of membranes shown in Figure 7B; arrowheads indicate specific proteins bands.
